# Proteomic Insights into Circadian Transcription Regulation: Novel E-box Interactors Revealed by Proximity Labeling

**DOI:** 10.1101/2024.04.18.590107

**Authors:** Manon Torres, Achim Kramer

## Abstract

Circadian clocks (∼24h) are responsible for daily physiological, metabolic and behavioral changes. Central to these oscillations is the regulation of gene transcription. Previous research has identified clock protein complexes that interact with the transcriptional machinery to orchestrate circadian transcription, but technological constraints have limited the identification of de novo proteins. Here we use a novel genomic locus-specific quantitative proteomics approach to provide a new perspective on time-of-day-dependent protein binding at a critical chromatin locus involved in circadian transcription - the E-box. Using proximity labeling proteomics at the E-box of the clock-controlled *Dbp* gene in mouse fibroblasts, we identified 69 proteins at this locus at the time of BMAL1 binding. This method successfully enriched BMAL1 as well as HDAC3 and HISTONE H2A.V/Z, known circadian regulators. New E-box proteins include the MINK1 kinase and the XPO7 transporter, whose depletion in human U-2 OS cells results in disrupted circadian rhythms suggesting a role in the circadian transcriptional machinery. Overall, our approach uncovers novel circadian modulators and provides a new strategy to obtain a complete temporal picture of circadian transcriptional regulation.

## Introduction

Circadian clocks are molecular oscillators that are present in most mammalian cells and operate on an approximate 24-hour cycle. They coordinate the daily rhythms of molecular, physiological and behavioral processes. These clocks play a central role in maintaining health, and disruptions caused by factors such as shift work and jet lag have been linked to the onset of various diseases such as cancer, diabetes, obesity, cardiovascular and neurodegenerative diseases. Therefore, deciphering the basic mechanisms that control circadian rhythms is essential for the development of effective health strategies.

At the heart of molecular oscillators is the regulation of circadian transcription. Several cis-regulatory proteins and enhancer elements, including E-box, Rev-ErbA/ROR response element (RRE) and D-box, have been identified as necessary elements of transcriptional-translational feedback loops (TTFLs) that drive cell-autonomous oscillations. The transcription factors BMAL1 and CLOCK form a protein complex that rhythmically binds to E-box enhancer elements typically located near the transcription start site of clock or clock-controlled genes (Gekakis et al., 1998; Ripperger et al., 2000; Ripperger and Schibler, 2006). This complex promotes the transcription of its own inhibitors, especially PERs and CRYs, which in turn form a repressor complex. This inhibitory complex attenuates CLOCK:BMAL1 transcriptional activity with a delay of several hours (Aryal et al., 2017; Gabriel et al., 2021). The oscillation is strengthened by REV-ERB and ROR proteins that bind to the RREs in the *Bmal1* promoter.

The network of clock-controlled genes directly regulated by CLOCK:BMAL1 includes several transcription factors and coregulators. For example, the CLOCK:BMAL1 complex orchestrates the rhythmic expression of the D-box binding PAR BZIP transcription factor (*Dbp*) gene (Ripperger et al., 2000), which in turn controls a large number of genes containing D-box sequences (Bozek et al., 2007; Yoshitane et al., 2019). This system allows for broad temporal control of downstream factors via rhythmic transcriptional regulation.

Mechanistically, the transcriptional activity of BMAL1:CLOCK is regulated at multiple levels, including its abundance in the nucleus (Kondratov et al., 2003; Kwon et al., 2006), formation of the BMAL1:CLOCK heterodimer, DNA binding and histone modifications (Asher et al., 2008; Beytebiere et al., 2019; Curtis et al., 2004; Etchegaray et al., 2003; Hirayama et al., 2007; Koike et al., 2012; Nakahata et al., 2008; Shi et al., 2016), as well as protein turnover. Heterodimerization and turnover of BMAL1 and CLOCK are closely related to post-translational modification processes, including phosphorylation (Aryal et al., 2017; Klemz et al., 2021; Kwak et al., 2013; Sahar et al., 2010; Spengler et al., 2009; Tamaru et al., 2009; Yoshitane et al., 2019), ubiquitylation (Kwon et al., 2006; Stratmann et al., 2012), or SUMOylation (Cardone et al., 2005; Lee et al., 2016). In addition, characterization of the BMAL1 and CLOCK cistromes in mouse liver has shown that DNA binding by the BMAL1:CLOCK complex does not determine the transcription phase of most target genes (Menet et al., 2012; Trott and Menet, 2018) highlighting the influence of additional transcriptional modulators on clock-regulated gene expression. This suggests that BMAL1:CLOCK regulation of clock-regulated gene expression relies on cooperation between CLOCK:BMAL1 and other transcription factors, allowing for interplay with other signaling pathways.

The study of proteins involved in the circadian regulation of transcription has so far relied on antibody-based chromatin immunoprecipitation (ChIP), protein-protein interaction (PPI) or cistrome analysis. However, these methods are only partially suitable for the de novo identification of proteins involved in the circadian transcription machinery. Recently, proximity labeling in combination with mass spectrometry proteomics has emerged as a complementary strategy (Niinae et al., 2021). Techniques based on proximity biotinylation have been used to profile various protein interactions, including insoluble protein complexes and temporal protein interaction profiling. Engineered peroxidases (e.g. APEX, APEX2, HRP) oxidize biotin-phenols to biotin-phenoxyl radicals in the presence of H_2_O_2_, and the resulting radicals biotinylate electron-rich amino acids (e.g. Tyr, Trp, Cys, His). Engineered peroxidases have been used to develop new tools to identify regulatory proteins involved in the control of gene expression. In their studies, Gao et al. (2018) and Myers et al. (2018) used APEX or APEX2 fused to a catalytically dead Cas9 mutant (dCas9) for the biotinylation of DNA-binding proteins near specific DNA elements (Gao et al., 2018; Myers et al., 2018).

Here, we adapted a DNA-protein proximity labeling method based on APEX2 biotinylation and subsequent liquid chromatography mass spectrometry (LC-MS/MS) proteomics to identify proteins that bind to the E-box region of the classical clock-controlled gene *Db*p in mouse NIH3T3 fibroblasts. We identified 69 proteins that are present at the E-box of the first intron of *Dbp* at the time of maximal BMAL1, including proteins already known to be involved in circadian transcription via E-boxes, such as BMAL1, HDAC3 and H2AZ, as well as previously unknown E-box proteins such as XPO7 and MINK1. The latter play a critical role in normal E-box-mediated transcription, as their depletion in human U-2 OS cells leads to disturbed circadian rhythms. This demonstrates that our proximity labeling approach is suitable for the unbiased identification of novel circadian transcription modulators.

## Results

### Protein-DNA proximity labeling to study circadian E-box regulators

To identify proteins localized at the E-box of *Dbp* at a given time point, we used a novel proximity labeling method, in which a deactivated CAS9 nuclease is fused to an engineered ascorbic acid peroxidase (APEX2) enzyme (dCAS9-APEX2). In this approach, CRISPR-CAS9 technology is used to guide the dCAS9-APEX2 fusion protein (CASPEX complex) to a desired location using specially designed sgRNAs. APEX2 is a peroxidase that catalyzes the biotinylation of surrounding proteins when biotin-phenol and H_2_O_2_ are added to the culture medium (**Figure 1A**). The biotinylated proteins can then be enriched due to their strong affinity for streptavidin and identified by proteomic methods such as immunodetection or mass spectrometry. This rapid labeling reaction (1 minute) is particularly advantageous to achieve the temporal resolution required in circadian studies.

**Figure 1:**
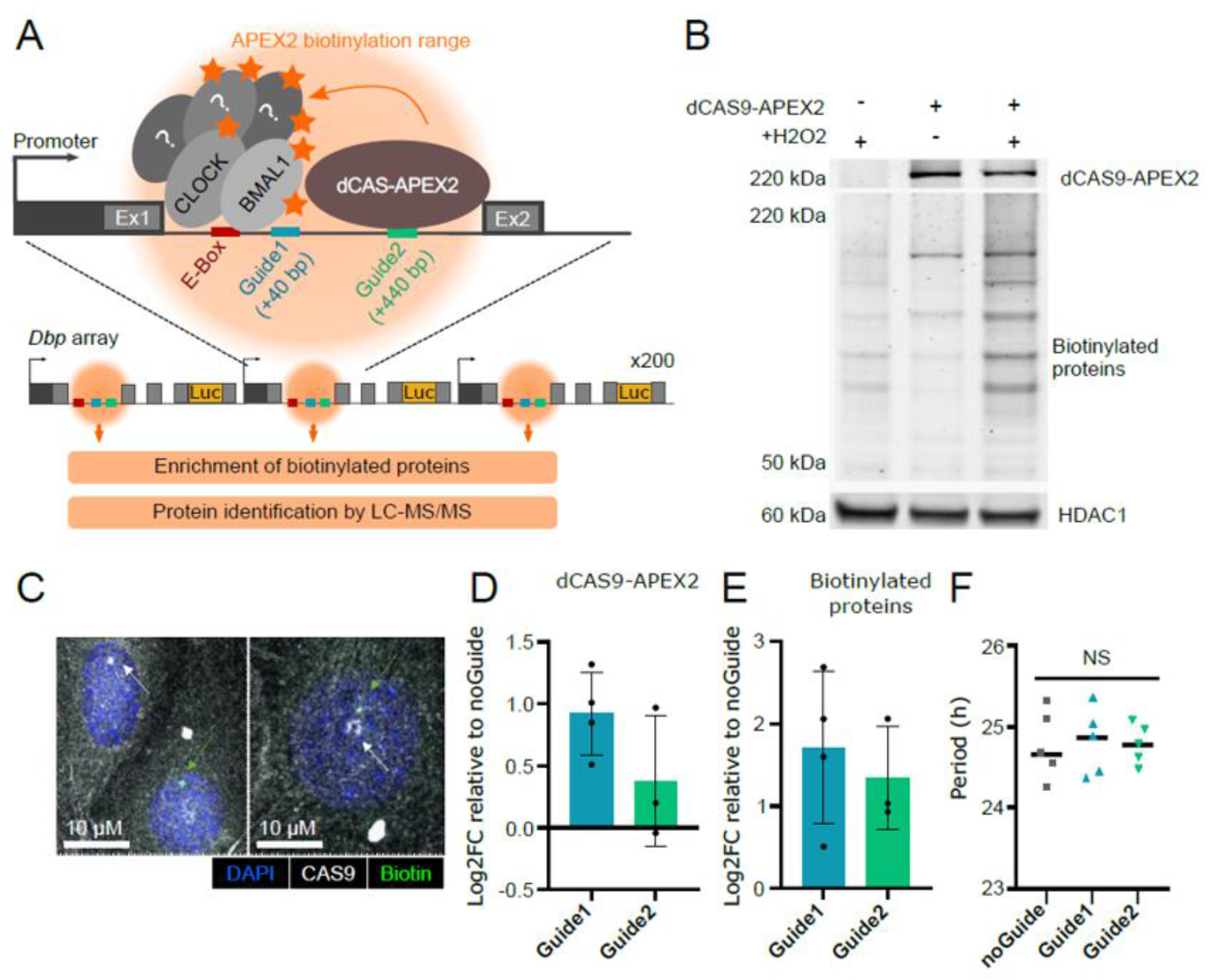
Proximity labelling of E-box proteins using the CASPEX method. **A**: Schematic description of the method. The sgRNAs target the the dCAS9-APEX2 fusion protein downstream of the E-box of the first intron of the Dbp gene. After induction, dCAS9-APEX2 binds the *Dbp* gene, and upon addition of biotin-phenol and H2O2, the APEX2 peroxidase induces biotinylation of proteins in close proximity. This reaction occurs simultaneously at the 200 repeats of the *Dbp* gene expressed in this Dbp(luc) array NIH3T3 mouse cell line. The biotinylated proteins are then purified using their affinity for streptavidin and subsequently identified by LC-MS/MS. **B:** Immunodetection of dCAS9-APEX2 and validation of its enzymatic activity. dCAS9-APEX2 was immunodetected in nuclear lysates of Dbp(luc) array NIH3T3 mouse cells using an anti-CAS9 antibody. In the same samples, biotinylated proteins were detected using horseradish peroxidase (HRP)-conjugated streptavidin followed by chemiluminescence imaging. All cells were treated with biotin-phenol, and H_2_O_2_ was added where indicated. **C:** Example of confocal microscopy fluorescence images of dCAS9-APEX2 (anti-CAS9 antibody) and biotinylated proteins (- Streptavidin, Alexa Fluor™ 514 conjugate) in Dbp(luc) array NIH3T3 mouse cells, expressing both dCAS9-APEX2 and guide sgRNAs, 24 hours after dCAS9-APEX2 induction and fixed after addition of APEX2 substrates (biotin-phenol and H_2_O_2_). High intensity aggregates are observed in the nucleus, similar to the BMAL1-YFP aggregates on the Dbp(luc) array in NIH3T3 cell model (Stratmann et al. 2012). **D** and **E:** Quantification of aggregates of dCAS9-APEX2 (**D**) and biotinylated proteins (**E**) from confocal microscopy images (as shown in **C**). For each field-of-view, the percentage of nuclei containing an aggregate of dCAS9-APEX2 or biotinylated proteins was quantified and expressed as log 2-fold-change compared to the control cell type (dCAS9-APEX2 + noGuide); n=3-4 biological replicates, each with 5-12 fields-of-view per cell line. **F:** Circadian period of different cell lines. Period values were extracted from 3 days bioluminescence time series, recorded 24h after synchronization and dCAS9-APEX2 induction, using the Chronostar software and did not show significant differences (one-way ANOVA, n=5, p>0.05, i.e. not significant (NS)).

To improve the enrichment of E-box-binding proteins, we used a Dbp(luc) array NIH3T3 mouse cell line, a creation of the Schibler lab (also characterized in our lab), which contains approximately 200 copies of the *Dbp* gene, a known E-box-regulated gene (Klemz et al., 2021; Stratmann et al., 2012). In previous studies, such tandem arrays were used to monitor the binding of fluorescently labeled transcription factors to DNA (Bosisio et al., 2006; Karpova et al., 2008; McNally et al., 2000; Stratmann et al., 2012). We selected this cell line to overcome potential sensitivity limitations associated with proximity labeling technology. Dbp(luc) array NIH3T3 cells were stably transfected with an inducible dCAS9-APEX2 construct, which allowed the induction of dCAS9-APEX2 expression by addition of doxycycline to the culture medium.

We first determined the concentration of doxycycline required for robust expression of dCAS9-APEX2 (0.25 µg/mL) in the nuclear fraction (**Supplemental Figure S1**). We then validated the enzymatic activity or biotinylation activity of dCAS9-APEX2. Using Western blot analysis, we assessed the amount of biotinylated proteins in the nuclear lysate as a function of the expression of the CASPEX complex and/or the addition of its substrates (biotin-phenol and H2O2). The presence of both dCAS9-APEX2 and the substrates resulted in a significant increase in the total intensity of biotinylated proteins (**Figure 1B, Supplemental Figure S2**).

To specifically direct the dCAS9-APEX2 fusion protein to the E-box in the first intron of the *Dbp* gene, we designed sgRNA guides targeting positions at different distances from the E-box. The sgRNA sequences were positioned within 500 bp of the targeted E-box to ensure effective biotinylation by the APEX2 peroxidase (**Figure 1A**). Next, we validated the ability of our sgRNA to guide the dCAS9-APEX2 complex to the *Dbp* array by quantifying both the accumulation of dCAS9-APEX2 and biotinylated proteins on the *Dbp* array. Biotinylation was induced in the cells 24 hours after CASPEX induction by doxycycline, followed by immediate fixation of the cells. The dCAS9-APEX2 complex was immunodetected with a CAS9-specific antibody, and biotinylated proteins were detected with streptavidin-conjugated fluorophores (**Figure 1C**). Confocal microscopy revealed fluorescent aggregates in Dbp(luc)-array NIH3T3 cells expressing the dCAS9-APEX2 complex. Cells expressing Guide 1 at +40 bp or Guide 2 at +440 bp showed an increased amount of CASPEX aggregates (1.9- and 1.3-fold change, respectively, **Figure 1D**) and a higher number of aggregates of biotinylated proteins (3.7- and 2.7-fold change, respectively, **Figure 1E**) compared to cells expressing a noGuide construct. These results indicate that these guides effectively bring dCAS9-APEX2 to the E-boxes of the *Dbp* array to induce biotinylation of the proteins localized on it.

To ensure that the presence of CASPEX near the E-box of *Dbp* does not interfere with the circadian machinery of rhythmic *Dbp* expression, we compared the rhythmicity of cells expressing Guide1 and Guide2 with cells expressing the noGuide construct. Since the Dbp(luc) array NIH3T3 cells were equipped with a luciferase reporter in the sequence of each *Dbp* gene repeat, we could monitor *Dbp* E-box-dependent transcription via live-cell bioluminescence recording. Cells expressing Guide1, Guide2, or noGuide were synchronized with dexamethasone, and bioluminescence signals were monitored for approximately 4 days. Circadian rhythmicity in cells expressing sgRNA guides directing CASPEX near the E-box of Dbp did not differ from the control, i.e. there were no significant changes in period length (**Figure 1F**) or other circadian parameters (**Supplemental Figure S3**).

### Identification of proteins located at the E-box of the Dbp gene

Our next goal was to identify the proteins located at the E-box of the *Dbp* gene. To this end, we synchronized the circadian rhythms of our cell lines with dexamethasone and 24 hours later, at the time of maximal BMAL1 binding to E-boxes (Stratmann et al., 2012), we induced biotinylation of proteins in the vicinity of dCAS9-APEX2 targeted close to the E-box of the *Dbp* gene. The biotinylated proteins were subsequently enriched from the nuclear lysates with streptavidin-coated magnetic beads and identified by label-free quantitative LC-MS/MS proteomics (**Figure 2A**). The experiment included cells expressing two different sgRNA guides (Guide1 and Guide2) and a control cell type transduced with an empty vector guide (noGuide). A total of 2687 proteins were detected in at least three replicates of at least one cell type (with a standard deviation between replicates <1). In Guide1, Guide2 and noGuide cell types we identified 2074, 2281 and 2105 proteins, respectively, which were detected in at least three replicates (**Figure 2B**, **Supplemental Table S1**) suggesting a substantial enrichment of non-specific proteins, possibly due to APEX2-independent biotinylation, high overall expression of dCAS9-APEX2 in the nucleus, or contamination by non-biotinylated proteins.

**Figure 2:**
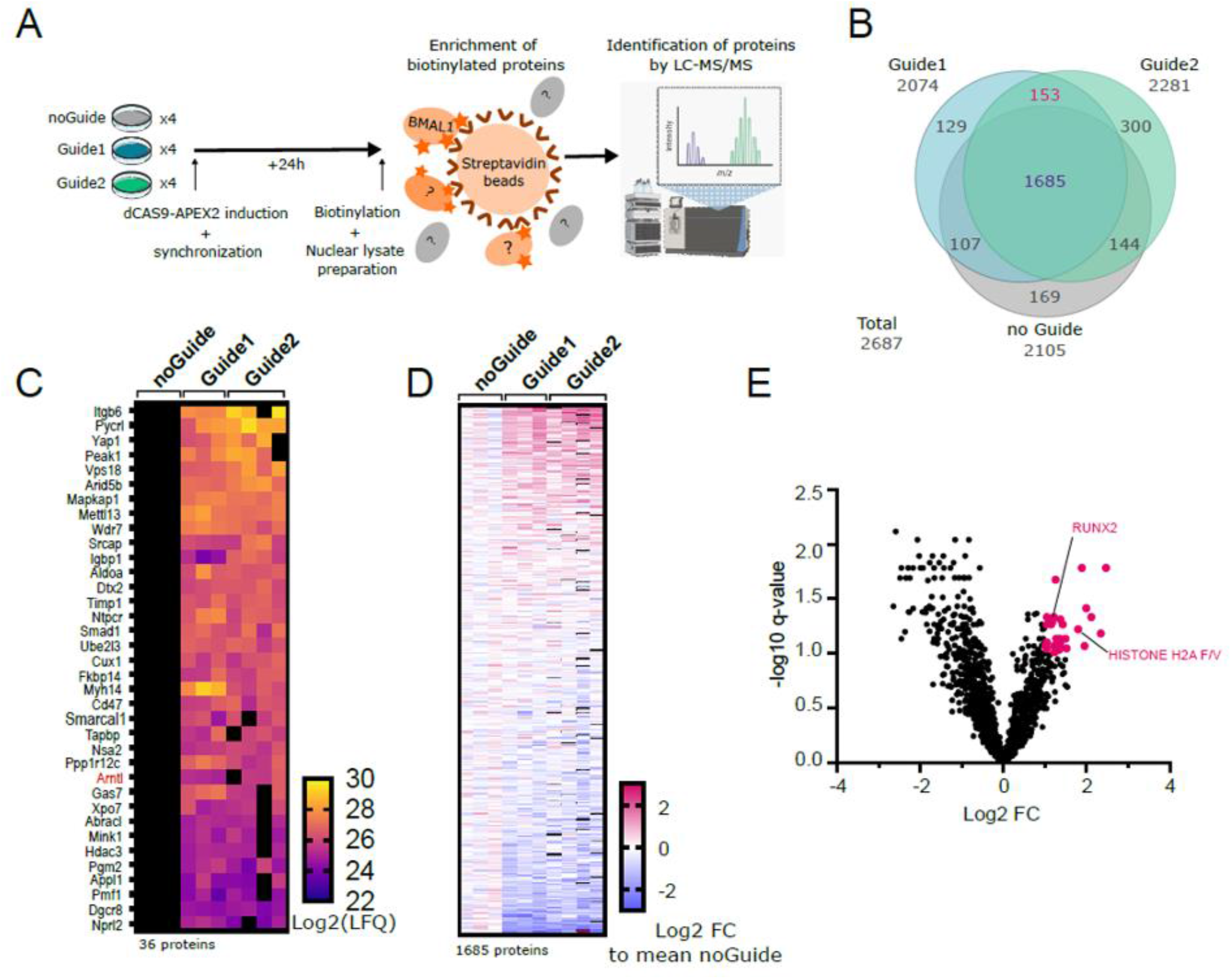
Identification of proteins located at the E-box of the *Dbp* gene using mass spectrometry. **A**: Experimental pipeline. **B**: Number of identified proteins in total, per cell type and overlap between cell types. Proteins were considered reliably identified if they were detected in ≥3 replicates of a cell type and the standard deviation between replicates of the same cell type was <1. **C**: Detection intensity (in Log2LFQ) of the 36 proteins that were only detected in Guide1 and Guide2 but not in noGuide cells (≥3 replicates per cell type). **D**: Fold change of the detection intensity of all samples compared to the mean intensity of noGuide for the 1685 proteins that were detected in all three cell types. Warm colors indicate a positive fold change compared to noGuide cells and cold colors indicate a negative fold change (≥3 replicates per cell type). **E**: Volcano plot of the log2 fold change of proteins versus the -log10 q-value in Guide1 and Guide2 compared to noGuide cells. The proteins with significant enrichment (i.e., fold change >2 and q-value <0.1) are highlighted (≥3 replicates per cell type).

To specifically identify proteins located on the E-box of *Dbp*, we required that a protein must be detected in both Guide1 and Guide2 cell types and either absent or significantly lower in the noGuide control. Using these criteria, we identified 36 proteins that were detected exclusively in Guide1 and Guide2 cells but not in noGuide cells (**Figure 2C, Supplemental Table S1**). We considered these proteins to be specific for CASPEX localization to the E-box of *Dbp*. In addition, for the 1685 proteins detected in all three cell types we analyzed of the fold change in intensity of each replicate compared to the mean intensity of the noGuide control and found that ∼60% of these proteins had lower levels in Guide1 and Guide2 compared to noGuide cells, while ∼40% showed enrichment in Guide1 and Guide2 compared to noGuide cells (**Figure 2D**). To identify additional E-box binding candidates, we used both their fold-change (≥2) and their statistical enrichment in the Guide compared to the noGuide cells using the false discovery rate (FDR) approach (q-value <0.1) as further criteria. This analysis identified 33 additional proteins (**Figure 2E, Supplemental Table S1**).

In total, we identified 69 proteins that are enriched in both Guide1 and Guide2 cells and are absent or significantly lower in the noGuide control cells. These proteins are presumably located near the E-box, and their role in the circadian transcriptional machinery can be further investigated.

### Proteins enriched at the E-box of the *Dbp* gene

Among the enriched proteins, we found the central transcription protein BMAL1, as expected for this time point (**Figure 3A**) demonstrating the specificity of the approach. In addition to BMAL1, our protein list included six other transcription factors or coactivators (YAP1, RUNX2, SMAD1, CUX1, SRCAP, and ARID5B (**Figure 3A**)) and three chromatin modifiers (HDAC3, HISTONE H2A.V/Z, and SMARCAL1 (**Figure 3B**)). Furthermore, we highlighted three proteins involved in posttranslational modifications, i.e. the kinase MINK1, the ubiquitin-conjugating enzyme UBE2L3 and the ubiquitin-protein ligase DTX2 (**Figure 3C**), as well as three proteins associated with protein transport (XPO7, APPL1, and MVP (**Figure 3D**)).

**Figure 3:**
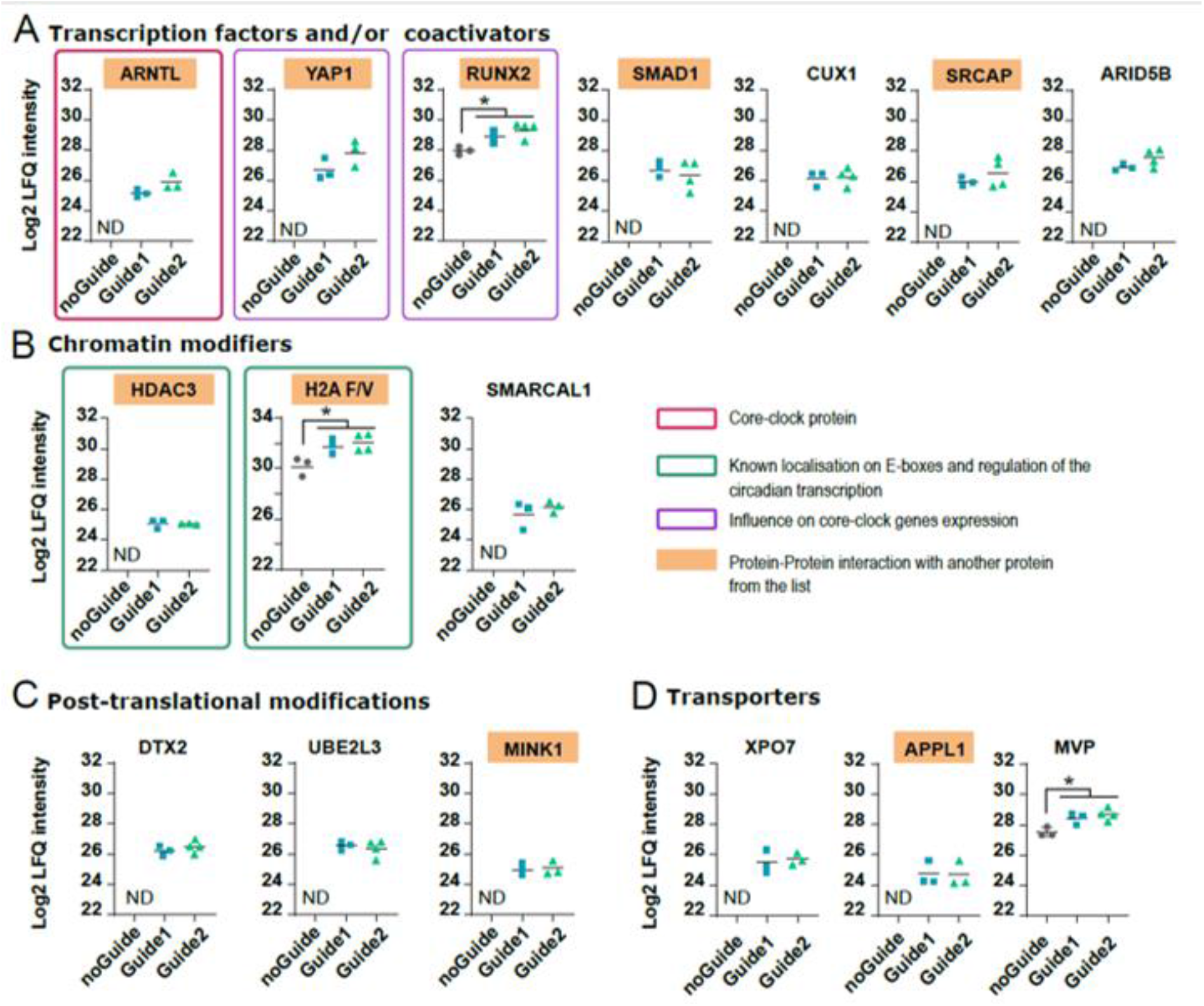
Identified proteins and their functional role Log2 LFQ intensity values of the proteins identified in Guide1 and Guide2 compared to noGuide control cells. ND= not detected; statistical significance is defined as fold change ≥2 and a q-value <0.1.

Interestingly, the detection of the histone variants H2A.Z and HDAC3 is consistent with their known roles in the circadian transcriptional machinery in liver cells (Tartour et al., 2022) (Shi et al., 2016), suggesting a mechanism that goes beyond liver specificity. In addition, both RUNX2 and YAP1 have been identified as transcription factors that influence circadian rhythmicity. In the mouse suprachiasmatic nucleus, knockdown of RUNX2 resulted in altered mPer2:luc rhythms in embryonic cultures, while Runx2^+/-^ mice exhibited a prolonged period of free-running activity (Reale et al., 2013). YAP1 is shown to repress clock gene expression in undifferentiated pleomorphic sarcomas, thereby promoting cell proliferation (Rivera-Reyes et al., 2018) and available ChIP-seq datasets from cancer cell lines suggest localization in several E-box-regulated core clock genes (Stein C et al., 2015; Zanconato et al., 2015). It remains to be determined whether any of these effects on circadian rhythmicity are mediated by involvement in the circadian transcriptional machinery in healthy cells.

In addition, for several of the identified proteins, we found reports of protein-protein interactions with other candidates from our list, such as RUNX2 with YAP1 and SRCAP (Guo et al., 2021; Thomsen et al., 2021) SMAD1 with YAP1 and MINK1 (Alarcón et al., 2009; Chen et al., 2022; Kaneko et al., 2011) HISTONE 2AZ with SRCAP (Liang et al., 2016; Tartour et al., 2022), and HDAC3 with APPL1 (Fan et al., 2019).

Overall, for many proteins identified using this approach, we found data in the literature that indeed suggest their localization at the E-box of Dbp and for their interaction with each other.

### New regulators of the circadian oscillator

To demonstrate the utility of this method in identifying novel proteins involved in the regulation of the circadian molecular oscillator, we investigated the effect of downregulating the expression of selected identified proteins on the oscillation of a *Bmal1*-luciferase reporter construct. To specifically characterize candidates relevant for the human circadian oscillator, we used the human osteosarcoma cell line U-2 OS as a well-characterized circadian oscillator model (Maier et al., 2021). We used multiple shRNAs for each target protein because of possible off-target effects that could produce false positive or false negative results. Among the identified proteins highlighted in Figure 3, we selected candidates for knockdown based on a preliminary screening of knockdown efficiency and normal cell growth after knockdown (not shown). Based on these criteria, we present the results for two candidate proteins: the nuclear export/import protein XPO7 and the kinase MINK1.

For XPO7, all four constructs resulted in a depletion of XPO7 transcript levels between 91 % and 95 % compared to the control (**Figure 4A**) and a shortening of the circadian period between 0.5 and 1.3 hours for three of the four shRNA constructs (**Figure 4B**). The amplitude-damping ratio was not substantially altered compared to the non-silencing control (**Figure 4C**). Similarly, we targeted the kinase MINK1 with four shRNA constructs, resulting in knockdown efficiencies between 87 % and 93 % (**Figure 4D**) and a shortening of the period between 0.7 and 1.3 hours for three of the four shRNA constructs (**Figure 4E**). Again, the amplitude-damping ratio was not substantially altered (**Figure 4F**).

**Figure 4:**
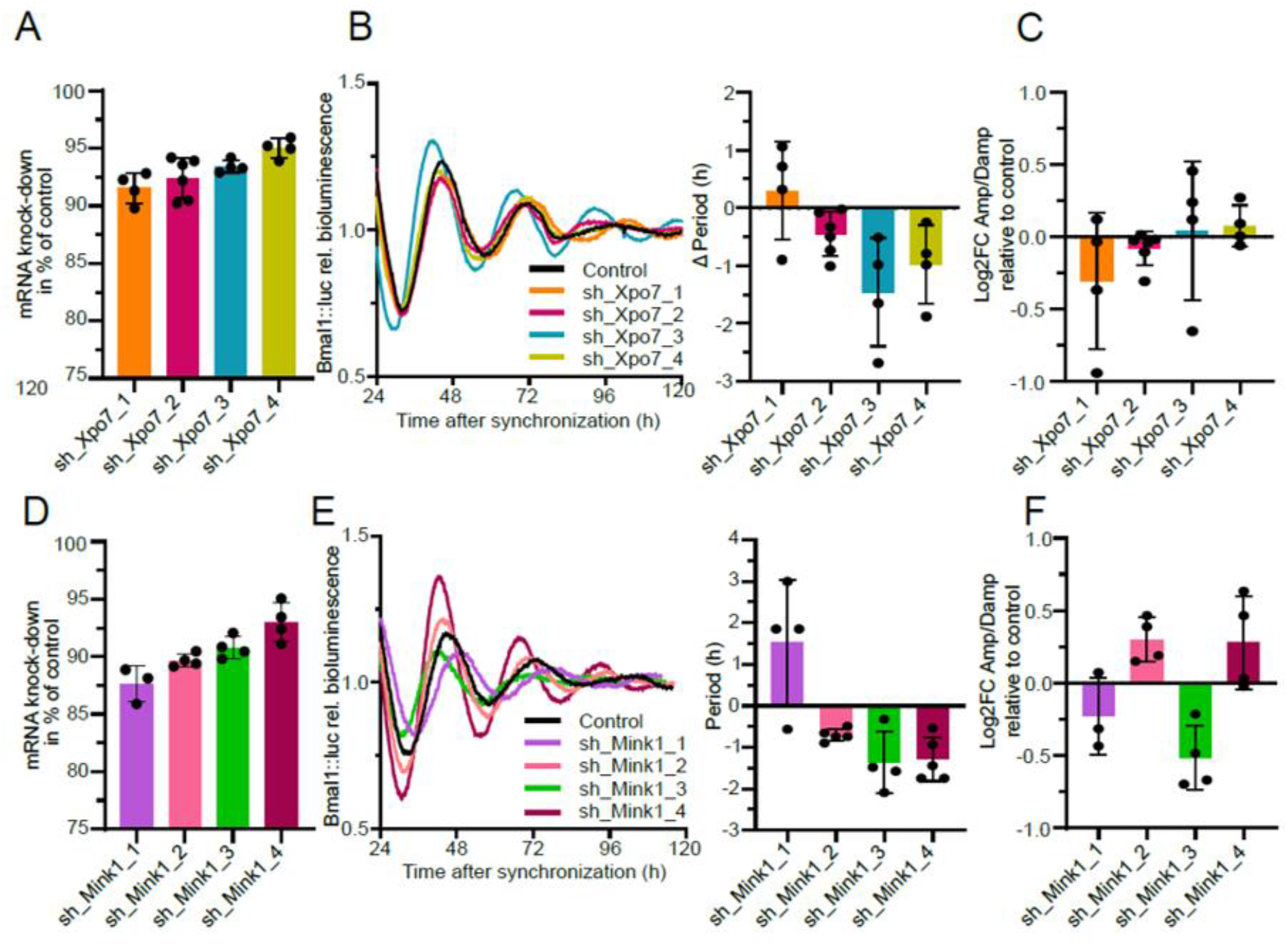
Depletion of identified E-box enriched proteins affects circadian dynamics. **A** and **D**: Quantification of knockdown efficiency by RT-qPCR 48 hours after synchronization (n=3 to 6 biological replicates, mean values ± SEM). **B** and **E**: Left panel: representative time series of *Bmal1*-luciferase U-2 OS cells transduced with indicated knockdown constructs. Right panel: quantification of changes in circadian period compared to non-silencing control (n=3 to 6 biological replicates, with at least three technical replicates, mean values ± SEM) **C** and **F**: Log2 fold change of amplitude-damping ratio after shRNA-mediated knockdown compared to non-silencing control (n=3 to 6 biological replicates, with at least three technical replicates, mean values ± SEM).

In summary, depletion of both XPO7 and MINK1 leads to a shortening of the period of the circadian oscillator. Since both proteins were identified to be enriched at the *Dbp* E-box, it is very likely that act on circadian dynamics via modulating circadian transcription.

## Discussion

The identification and characterization of proteins involved in the circadian transcriptional machinery on key cis-regulatory elements depends on the ability to selectively enrich proteins that are in close proximity to a particular genomic sequence at a given time point. Previous methods aiming at this task relied on chemical cross-linking to immobilize proteins in their spatial and temporal context. However, chemical cross-linking leads to changes in the protein structure that can interfere with the identification by mass spectrometry (MS) (Toews et al., 2008). Here, we use the biotinylating enzyme APEX2 targeted close to the E-box of the *Dbp* gene by a catalytically inactive CAS9 and specifically designed sgRNAs. In contrast to previous approaches, this method does not require chemical cross-linking and does not require the availability of specific antibodies. In addition, the reaction time of APEX2 of only one minute allows labeling of proximal proteins with a high time resolution. This unbiased characterization of the interactome at a specific chromosomal locus complements the chromatin immunoprecipitation (ChIP)-seq method, which delineates the multiple binding regions of a particular protein of interest.

In this first study, we focused on the E-box of the *Dbp* gene, a known E-box regulated gene whose binding to core clock proteins has previously been characterized by ChIP-seq strategies. To increase the enrichment efficiency, we used a *Dbp* array in the mouse fibroblast NIH3T3 cell line, which expresses 200 copies of the *Dbp* gene as tandem repeats. After establishing stable expression of dCAS9-APEX2 and the sgRNAs in this cell model, we observed, as expected, substantial accumulation of the fusion protein on the *Dbp* array, but also in other cell compartments (especially in the cytoplasm, not shown), leading to non-specific biotinylation of proteins. To distinguish E-box proteins from non-specific biotinylated proteins, we used two cell lines with different sgRNAs for chromosomal targeting, resulting in overlapping labeling radii. Using our criteria, we identified a total of 69 proteins that are most likely enriched in the E-box of the *Dbp* gene.

Using the extensive literature on E-box binding proteins, we found evidence for localization at the E-box of *Dbp* for several of the proteins identified here. The key proteins expected at the E-box of *Dbp* at this specific time point are BMAL1 and CLOCK. Original work on the *Dbp* array NIH3T3 cell line showed that overexpressed BMAL1-YFP reaches its highest binding to the *Dbp* gene array approximately 24 hours after cell synchronization by dexamethasone (Stratmann et al., 2012). Accordingly, using the same cell line and inducing proximity labeling 24 hours after cell synchronization, we enriched endogenous BMAL1 specifically in both cell types where the labeling enzyme is brought in proximity to the E-box of *Dbp*. In ChIP-seq time series experiments using mouse liver, both BMAL1 and CLOCK were detected at the E-box of the first intron of *Dbp* at the time of highest BMAL1 binding (Koike et al., 2012). However, in our data set, CLOCK was not enriched in any sample of the cell lines analyzed, including in the noGuide control, indicating that the amount of biotinylated endogenous CLOCK was below our detection limit. In addition to the core clock protein CLOCK and BMAL1, it has been described that p300 interacts with CLOCK-BMAL1 at E-boxes during this phase of circadian transcription (Koike et al., 2012). In our data set, p300 is robustly detected in all cell types, including the noGuide control, but was not significantly enriched in cells expressing E-box-specific sgRNA (**SupplementalTable S1**). This could mean that p300 is not specifically localized to the E-box of the first intron of Dbp, which is consistent with the p300 ChIP-seq results from mouse liver. However, it could also be an artifact resulting from the inverse relationship between protein abundance and the ability to determine when it is enriched over background.

Similarly, six of the RNA polymerase II subunits were detected with high intensity in all cell types and replicates, with no statistical enrichment in the guide cell lines, likely reflecting their overall abundance. The H2A.Z variant histone 1 and the isoform H2A.Z variant histone 2 were found significantly enriched in our sgRNA cells compared to noGuide cells. This is consistent with H2A.Z ChIP-seq data showing that in mouse liver H2A.Z not only flanks the core clock binding sites of E-box regulated genes such as *Per2* or *Cry1*, but is also required for the binding of BMAL1 to E-box controlled genes, including *Per1*, *Per2* and *Dbp* (Tartour et al., 2022). Furthermore, the same study suggests that H2A.Z nucleosome incorporation may be regulated by SRCAP and/or the p400-TIP60 chromatin remodeler complexes, which agrees with the specific enrichment of SRCAP at the E-box of Dbp in our dataset.

HDAC3, a histone deacetylase enzyme, was identified exclusively in both sgRNA cell types and not in the noGuide control. In liver and muscle tissue, HDAC3 has been detected by ChIP-seq at many ROREs sites together with REV-ERBα, but also at many E-box-regulated genes (Feng et al., 2011; Hong et al., 2017). Although HDAC3 has been shown to be recruited to the repressive REV-ERBα complex, it has also been described to associate with CLOCK and BMAL1 in the nucleus (Hosoda et al., 2009; Shi et al., 2016) and to promote the binding of BMAL1 to E-boxes in the main binding phase, independent of its enzymatic deacetylase activity (Shi et al., 2016).

Overall, our study represents a novel strategy to characterize the landscape of the circadian cistrome and to identify novel regulators of the complex machinery of circadian transcription in an unbiased manner. With this approach, not only known E-box regulators can be identified, but also new players in the circadian transcription machinery can be discovered at other chromatin regions. Several of the proteins identified here have not yet been described as components of the circadian machinery. Among them, we have identified two proteins that are required for normal circadian dynamics: the transporter protein XPO7 and the kinase MINK1. Our study is therefore an excellent starting point not only to discover new circadian modulators, but also to characterize the chromatin binding dynamics of these regulators and thus obtain a complete time-resolved picture of circadian transcriptional regulation.

## Material and methods

### Cell lines and culture

Mouse fibroblast NIH3T3 cells stably expressing a tandem array of ∼250 *Dbp* copies (Dbp(luc) array NIH3T3) were a kind gift from Ueli Schibler (University of Geneva) (Stratmann et al., 2012). Exon 4 of the *Dbp* gene contains an open reading frame for luciferase (luc) preceded by an internal ribosome entry site (IRES) to allow monitoring of transgene expression on the array by luminescence recording.

Human osteosarcoma U-2 OS *Bmal1*-luciferase cells (ATCC HTB-96) stably expressing firefly luciferase from a 0.9-kb mouse *Bmal1* promoter fragment (kind gift from Steven Brown, University of Zurich). All cell types were grown in DMEM medium with 10 % fetal bovine serum, 25 mM HEPES and antibiotics (100 U/mL penicillin, 100 μg/mL streptomycin).

### Generation of the Dbp(luc) array NIH3T3 dCAS9-APEX2 cell line

The inducible CASPEX expression plasmid (Addgene #97421) was stably transfected into Dbp(luc) array NIH3T3 cells using Lipofectamine™ 3000 from Invitrogen. Non-transduced cells were eliminated by both antibiotic selection (puromycin 10 μg/mL) and fluorescence-activated cell sorting (FACS). The inducible expression of dCAS9-APEX2 after doxycycline treatment was confirmed by SDS-PAGE and Western blot. The same cell line was used for all subsequent experiments.

### Design and cloning of guide RNAs

Guide RNA sequences were generated using the CRISPOR web tool (Concordet and Haeussler, 2018) and designed to target the E-box of the first intron of the *Dbp* gene with a maximum distance of 500 bp. Sequences with a high on-target score were cloned into the lentiGuide-Hygro-dTomato (Addgene #99376), following the guidelines of (Shalem et al., 2014).

### Production and transduction of lentiviruses

Lentiviruses were produced in HEK293T cells as described in (Maier et al., 2021). The virus-containing supernatants were filtered and the target cells were transduced with the virus filtrate. Dbp(luc) array NIH3T3 dCAS9-APEX2 cells were stably transduced with guide RNA-containing lentiviruses. Non-transduced cells were eliminated by antibiotic selection (0.25 mg/mL hygromycin) as well as by FACS. U-2 OS *Bmal1*-luciferase cells were transduced with a lentivirus containing shRNA transduced and selected for 3 days (puromycin 10 μg/mL).

### APEX2-mediated labelling

Prior to protein labeling, circadian rhythms were synchronized by incubating cells with 1 μM dexamethasone (Sigma-Aldrich™, #D4902) for 20 minutes. They were then rinsed twice with warm PBS and normal culture medium containing 0.25 µg/ml doxycycline was added to induce CASPEX expression (Clontech #631311). 24 hours later, biotin-tyramide-phenol (Iris Biotech GNBH, #41994-02-9), resuspended in DMSO, was added to the culture medium at a final concentration of 500 μM. After incubation for 30 minutes at 37 °C, biotinylation of the proteins was induced by adding 1 mM H_2_O_2_ for exactly 60 seconds. After 60 seconds, the medium was aspirated and the cells were washed three times with ice-cold quenching buffer that inactivates APEX2 activity (TBS with 1 mM CaCl_2_, 10 mM sodium ascorbate, 1 mM sodium azide, 1 mM Trolox, 1 mM DTT).

### Cell lysis and enrichment of nuclear proteins

Nuclear proteins were isolated according to the Abcam nuclear fractionation protocol. On ice, cell medium was replaced with fractionation buffer (10 mM HEPES pH7.4, 1.5 mM MgCl_2_, 10 mM KCl, 0.5 mM DTT, 0.05 % NP40, 1:100 protease inhibitor cocktail) and cells were scraped, collected and incubated on ice for 10 min. After 10 min centrifugation at 4 °C and 720 g, the supernatant containing cytoplasm, membranes and mitochondria was aspired and discarded. The pellet containing the nuclei was suspended in a lysis buffer (5 mM HEPES pH 7.4, 1.5 mM MgCl_2_, 0.2 mM EDTA, 0.5 mM DTT, 26 % glycerol, 1:100 protease inhibitor cocktail) containing 300 mM NaCl to lyse the membranes and solubilize the DNA. Samples were homogenized with a 27G needle and incubated on ice for 30 minutes. After centrifugation at 24,000 g for 20 minutes at 4 °C, the supernatants containing nuclear proteins were separated. The protein concentration was determined using a BCA assay.

### SDS-PAGE and Western Blot

Equal amounts of proteins (50 to 100 µg) were denatured by heating for 5 minutes at 95 °C in 1:4 SDS loading buffer. Protein separation was performed by SDS-PAGE. Proteins were transferred to a nitrocellulose membrane overnight in 5 % methanol blotting buffer. Protein smears were visualized by incubation in Ponceau for 10 seconds. To reduce non-specific reactions, the membranes were incubated for 1 hour in TBS-T containing 5 % bovine serum albumin (BSA). The membranes were then incubated overnight with primary antibodies in TBS-T (αCAS9 1:1000, αHDAC1 1:1000), washed and then probed with the corresponding HRP-conjugated secondary antibodies (1:1000). Biotinylated proteins were detected by incubating the membranes with HRP-conjugated streptavidin (1:10 000). A chemiluminescence reaction was performed for protein visualization.

### Enrichment of biotinylated proteins for LC-MS/MS analysis

Twenty 20 cm dishes with confluent cells were used for each replicate of the proteomic experiments to ensure an input of 5 mg of nuclear proteins for each sample. To each sample, 500 µL of Dynabeads™ MyOne™ Streptavidin T1 (Invitrogen #65601) was added, followed by overnight incubation at 4 °C on a rotating wheel. Using a magnetic support, the streptavidin beads (and attached biotinylated proteins) were washed twice with cold lysis buffer, once with cold 1 M KCl, once with 100 mM Na_3_CO_2_ and twice in cold 2 M urea in 50 mM ammonium bicarbonate (ABC) (as described in (Myers et al., 2018)). The washing solution was removed and the proteins on the beads were resuspended in 20 μL urea buffer (2 M urea, 50 mM ammonium bicarbonate, pH 8.0), reduced in 12 mM dithiothreitol at 25 °C for 30 min and then alkylated with 40 mM chloroacetamide at 25 °C for 20 min. The samples were digested with 0.5 μg trypsin/LysC overnight at 30 °C. The peptide-containing supernatant was collected and the digestion was stopped with 1 % formic acid. The peptides were desalted and purified according to the stage-tip protocol (Rappsilber et al., 2003). Samples were eluted with 80 % acetonitrile/0.1 % formic acid, dried with Speedvac, resuspended in 3 % acetonitrile/0.1 % formic acid and analyzed by LC-MS/MS.

### Liquid chromatography-mass spectrometry

Peptides were analyzed on a reversed-phase column, a 20 cm long fritless silica microcolumn with an inner diameter of 75 μm, packed with ReproSil-Pur C18-AQ 3 μm resin (Dr. Maisch GmbH) using a 90-minute gradient at a flow rate of 250 nL/min with increasing concentration of Buffer B (from 2 % to 60 %) on a high performance liquid chromatography (HPLC) system (Thermo Fisher Scientific), ionized with an electrospray ionization (ESI) source (Thermo Fisher Scientific) and analyzed on a Thermo Q Exactive Plus instrument. The instrument was operated in data-dependent mode, selecting the 10 most intense ions in the MS full scans (ranging from 350 to 2000 m/z) with a resolution of 70 K, an ion count of 3×10^6^ and an injection time of 50 ms. Tandem MS was performed with a resolution of 17.5 K. The MS2 ion counting target was set to 5×10^4^ with a maximum injection time of 250 ms. Only precursors with charge state 2-6 were selected for MS2. The dynamic exclusion time was set to 30 s with a 10 ppm tolerance for the selected precursor and its isotopes. Raw data were analyzed with the MaxQuant software package (v1.6.3.4, PMID: 27809316) using the UniProt mouse database (MOUSE.2019-07) and the APEX2 construct sequence database with forward and reverse sequences. The search included variable modifications such as methionine oxidation, N-terminal acetylation as well as asparagine and glutamine deamidation. Carbamidomethylation of cysteine was defined as a fixed modification. The minimum peptide length was set at seven amino acids and a maximum of 3 failed cleavages were allowed. The FDR (false discovery rate) was set at 1 % for peptide identification. Unique and Razor peptides were considered for quantification. Retention times were recalibrated based on the integrated nonlinear time scaling algorithm. MS2 identifications were transferred between runs using the “Match between runs” option, with the maximum retention time window set to 0.7 min. The LFQ algorithm (label-free quantification) was activated.

### Immunofluorescence microscopy

Prior to APEX2-mediated labeling, cells were seeded to 90 % confluence on glass bottom chambers (Ibidi #80826) coated with fibronectin. At the end of the labeling protocol, the medium was removed and the cells were fixed by incubation in 4 % paraformaldehyde in PBS pH 7.4 for 10 minutes. After washing three times with cold PBS, cells were permeabilized by incubating three times for 5 minutes in Triton X-100 0.1 % in PBS alternating with 5-minute washes with cold PBS. dCAS9-APEX2 proteins were detected with a first antibody against CAS9 (1:1000) and a secondary antibody combined with fluorophores (10 µg/mL). Biotinylated proteins were detected by incubation with Streptavidin, Alexa Fluor™ 514 conjugate (5 µg/mL). Cell nuclei were labeled by DAPI staining (300 nM). Images were captured with a Nikon Scanning Confocal A1Rsi+ at 60X. The number of nuclei that had dCAS9-APEX2 or biotinylated protein dots was manually determined using Fiji (Schindelin et al., 2012) and expressed as a percentage of the total number of nuclei on the image.

### Bioluminescence imaging

Cells were seeded either on a white 96-well plate or in 35-mm NUNC dishes at the density required for confluence 1 to 3 days later. Circadian rhythms were then synchronized with 1 μM dexamethasone for 30 min, and cells washed with warm PBS, and placed in phenolred-free DMEM culture medium supplemented with 10 % fetal bovine serum, antibiotics (100 U/mL penicillin, 100 μg/mL streptomycin), and 250 μM D-luciferin (Biothema). Bioluminescence recordings were performed at 35 °C to 37 °C in a 96-well plate luminometer (TopCount, PerkinElmer) or LumiCycle (Actimetrics). The data were analyzed with the ChronoStar software (Maier et al., 2021). Briefly, the first 12 hours of recording were excluded, time series data were trend-eliminated by dividing by a 24-hour average, and rhythm parameters were estimated by fitting a cosine wave function with an exponential decay term.

### Determination of knockdown efficiency

Total RNA was purified using the Pure Link RNA Mini Kit (Thermo Fisher Scientific, #12183025) according to the manufacturer’s instructions with an additional on-column deoxyribonuclease I digestion. For quantitative reverse transcription PCR (RT-qPCR), RNA was reverse transcribed using the Moloney Murine Leukemia Virus Reverse Transcriptase Kit (Thermo Fisher Scientific, #28025013) with random hexamers (Thermo Fisher Scientific, #N8080127) according to the manufacturer’s instructions. RT-qPCR was performed using the SYBR Green Master Mix (Thermo Fisher Scientific, #A46112) and a CFX96 PCR system with CFX Manager software (Bio-Rad).

### Data analysis

We used of the Perseus software (Tyanova et al., 2016) to reduce our matrix of label-free quantification (LFQ) values by filtering out reversed proteins, proteins identified only by site, and contaminants. The values were converted to log2(x) values. We filtered out proteins that were not detected in at least three replicates of any of the cell types and had a standard deviation between replicates greater than 1. For proteins that were detected in all three cell types, we determined the statistical enrichment in Guide1 or Guide2 cells compared to the noGuide control using a multiple t-test with the false discovery rate approach (Benjamini-Hochberg method). Significant enrichment of a protein at the E-box position was defined by a p-value of less than 0.05 in and both Guide cell types compared to the control (defined by an unpaired t-test) and a fold-change greater than 2 in both cell types compared to the control.

## Acknowledgments

We thank all current and former members of the Kramer lab for technical and intellectual support. We thank the Ueli Schibler laboratory for the Dbp(luc) array NIH3T3 cells. We acknowledge the support of the FCCF at the German Rheumatism Research Center. We thank the Advanced Medical Bioimaging Core Facility (AMBIO) of the Charité for support in the acquisition of the imaging data. We thank Dr. Philipp Mertins and Dr. Marie-Luise Kirchner of the Core Unit Proteomics of the Berlin Institute of Health for their support in the design, processing and data acquisition of the liquid chromatography-mass spectrometry experiments. This work was funded by the Marie Skłodowska-Curie Individual Fellowships (846016 — CLOCK) granted by the Research Executive Agency (REA) under the mandate of the European Commission and the Deutsche Forschungsgemeinschaft (DFG, German Research Foundation) (KR1989/13-1).

## Author Contributions

Study design and conceptualization: A.K., M.T. Methodology: A.K., M.T. Investigation: M.T. Writing (original draft and editing): M.T. and A.K. Funding acquisition: M.T. and A.K.

